# Ultraviolet light degrades the mechanical and structural properties of human stratum corneum

**DOI:** 10.1101/614602

**Authors:** Z.W. Lipsky, G. K. German

**Affiliations:** Binghamton University

## Abstract

Prolonged exposure of human skin to sunlight causes photodamage, which induces the early onset of wrinkles and increased tissue fragility. While solar ultraviolet (UV) light is considered to have the most damaging effect, the UV range that is most harmful remains a topic of significant debate. In this study, we take a first step towards elucidating biomechanical photoageing effects by quantifying how exposure to different UV ranges and dosages impacts the mechanical and structural properties of human stratum corneum (SC), the most superficial skin layer. Mechanical testing reveals that irradiation of isolated human SC to UVA (365 nm), UVB (302 nm), or UVC (265 nm) light with dosages of up to 4000 J/cm^2^ notably alters the elastic modulus, fracture stress, fracture strain, and work of fracture. For equivalent incident dosages, UVC degrades SC the greatest. However, upon discounting reflected and transmitted components of the incident light, a generalized scaling law relating the photonic energy absorbed by the SC to the energy cost of tissue fracture emerges. This relationship indicates that no one UV range is more damaging than another. Rather, the magnitude of absorbed UV energy governs the degradation of tissue mechanical integrity. Subsequent structural studies are performed to elucidate the cause of this mechanical degradation. UV absorption scales with the spatial dispersion of desmoglein 1 (Dsg 1), a component of corneocyte cell-cell junctions, away from intercellular sites. Combining both scaling laws, we establish a mechanical-structural model capable of predicting UV induced tissue mechanical integrity from Dsg 1 dispersion.

**Statement of Significance:** Photoageing from the sun can produce early onset of skin wrinkles and an increase in tissue fragility that heightens the risk of rupture. While solar ultraviolet (UV) light is considered to have the most damaging effect, the UV range that is most harmful remains a topic of significant debate. In this study, we elucidate photoageing effects by quantifying how exposure to different UV ranges and dosages impacts the mechanical and structural properties of human stratum corneum (SC), the most superficial skin layer. Results establish a mechanical-structural model that relates the amount of UV energy absorbed by the tissue, irrespective of UV range, to the energy cost of tissue fracture and spatial dispersion of desmoglein 1.

## 1. INTRODUCTION

Solar radiation is comprised of wavelengths ranging between 220 (ultraviolet (UV)) and 3200 nm (infrared)(1). Due to its greater photon energy, the impact of UV radiation on skin is the most commonly examined, however studies have explored the effects of both visible and infrared radiation(2–4). UV light exposure has been linked with numerous health benefits such as increased vitamin D production and phototherapy of skin conditions such as eczema(5, 6). However, the health risks of UV overexposure are far more pervasive. UV exposure is associated with accelerated skin ageing(7), sunburn(8), and the promotion of skin cancers such as basel cell carcinomas, squamous cell carcinomas, and malignant melanomas(9). UV induced skin damage can be directly caused by DNA absorption of UV light, resulting in the formation of photolesions such as cyclobutane pyrimidine dimers or (6,4) photoproducts(8, 10–13), or indirectly through the generation of reactive oxygen species, which create oxidative stresses that damage DNA, proteins, and lipids(8, 14, 15). The range of UV light exposure can also influence the method and level of skin damage(11).

UV light is broken up into four distinct ranges: UVA (315-400 nm), UVB (280-315 nm), UVC(200-280 nm) and vacuum-UV (100-200 nm). Sunlight is comprised only of UVA and UVB light; UVC and vacuum-UV are fully attenuated by the atmosphere(16). However, exposure of skin to UVC light can occur when using welding equipment, sunbeds, or working within biosafety cabinets(17–19). In terms of overall damaging effects on full thickness skin, the World Health Organization defines UVC as the most damaging(20). However, different UV ranges do not impact skin tissue equally; UV light has wavelength-dependent penetration depths. While longer wavelength UVA can reach deep into the dermis, UVB and UVC light are primarily absorbed by the epidermis(21). As such, consideration for which tissue layer is affected needs to be accounted for. Moreover, reports of skin damage from different UV light ranges is evaluated primarily from a biological standpoint. Here damage is characterized by its impact on cellular DNA(13, 15, 22–24) and carcinogensis(25–28), rather than the potential for UV light to physically degrade skin integrity. While studies have explored UV light induced changes in skin elasticity(29, 30), and UVB’s effect on the mechanical properties of the most superficial skin layer, the stratum corneum (SC)(31), the relative impact of the different UV ranges on the structural and mechanical integrity of human skin remains unclear, as does the mechanism of degredation. In this study, we provide initial insight into the comparative damaging effects of different UV ranges by exploring how various UVA, UVB, and UVC light dosages change the mechanical and structural properties of human SC.

## 2. MATERIALS AND METHODS

### 2.1 Stratum corneum isolation

Full thickness 36 yrs. female breast skin specimens (*n* = 3) were received from Yale Pathology Tissue Services (New Haven, CT) within 24 hr. of elective surgery. Breast tissue was chosen because it is typically exposed only to low levels of solar UV exposure. An exempt approval (3002-13) was obtained to perform research using de-identified tissue samples pursuant to the Department of Health and Human Services (DHHS) regulations, 45 CFR 46.101:b:4. SC was isolated using a standard heat bath and trypsin technique(32–34). Once isolated, SC sheets were placed on plastic mesh, rinsed in deionized water, and dried for 48 hr. at room temperature and humidity. For uniaxial mechanical testing, SC samples were cut to a uniform 7 × 20 mm size (*n* = 142 total samples used for the study). For immunostaining studies, samples were cut to a uniform 3 mm diameter circle (*n* = 26 total samples used for the study). While full thickness skin exhibits mechanical anisotropy (35, 36), SC tissue exhibits isotropic mechanical properties(37, 38). As such, the orientation of SC samples cut from the isolated sheet are unimportant.

### 2.2 Ultraviolet Irradiation

UV irradiation of SC samples was performed using a UV Lamp (8 Watt EL Series, Analytik Jena US LLC, Upland, CA) with stand. For mechanical testing, SC samples were exposed to UVC (265 nm narrowband), UVB (302 nm narrowband) or UVA (365 nm narrowband) light for periods equating to incident light dosages of 10, 100, 200, 400, 800 and 4000 J/cm^2^. For structural studies, SC samples were exposed to UVC, UVB or UVA light for periods equating to incident light dosages of 100, 200, 400, and 800 J/cm^2^. For samples positioned 76 mm from the light source, the average intensities of the UVA, UVB, and UVC lamp bulbs are 1.5 mW/cm^2^, 1.6 mW/cm^2^, and 1.8 mW/cm^2^ respectively (39). To achieve equal dosages across the three UV ranges, exposure times were varied. For instance, a dosage of 10 J/cm^2^ required an exposure time of 111 min for UVA, 104 min for UVB, and 92 min for UVC. SC samples exposed only to ambient UV light were used for controls.

### 2.3 Mechanical characterization

After UV irradiation, samples were equilibrated for 24 hr. to either 25 or 100% relative humidity (RH) before testing. Equilibration to low or high RH conditions were achieved by placing specimens respectively in an airtight container filled with desiccant (Drierite 10-2 mesh, W.A. Hammond Drierite Company, Xenia, OH) or a hydration cabinet (F42072-1000, Secador, Wayne, NJ) with a base filled with deionized water. In both cases, RH conditions were monitored throughout the equilibration period using a hygrometer with probe (445815, Extech Instruments, Nashua, NH)). After equilibration, the mechanical properties of samples were evaluated using a uniaxial tensometer (UStretch, CellScale, Waterloo, ON, Canada) equipped with a 4.4 N load cell. The ends of each SC sample were taped to prevent slippage of the sample in the tensometer grips, leaving an exposed area of 7 × 10 mm. Individual SC samples were mounted into opposing tensometer grips, initially separated by 10 mm. Samples were strained until rupture at a constant strain rate of 0.012 s^−1^; similar to rates used in previous mechanical studies of skin(35). Tensile forces and grip separation were recorded at a frequency of 5 Hz. After mechanical testing, the average thickness of the ruptured SC sample was quantified using optical microscopy. Optical thickness measurements were taken a distance from the crack interface to prevent measuring reduced thicknesses arising from plastic deformation. Combinations of sample dimensions, and recorded force-displacement data were then used to derive engineering stress-strain curves, from which the average elastic modulus, fracture stress, fracture strain and work of fracture were extracted (3 ≤ *n* ≤ 9 independent samples for each UV range, dosage, and humidity).

### 2.4 Desmoglein 1 dispersion analysis

Irradiated and control samples (3 mm diameter circle, *n* = 2 for each condition) were first incubated with an anti-demoglein 1 (Dsg1) mouse monoclonal antibody (651110, Progen, Heidelberg, Germany) at a 1:500 ratio antibody to phosphate-buffered saline (PBS) at 4°C overnight (40, 41). Samples were then washed three times with PBS before being incubated in a 1:200 ratio of Alexa-Fluor 488 labeled goat anti-mouse IgG antibody (A-11029, Introgen, Carlsbad, CA) to PBS for 1 hr. at room temperature. Samples were then rewashed in PBS three times before being placed on a glass coverslip. Images of control and irradiated SC were recorded using a confocal microscope (Leica SP5, Wetzlar, Germany) and 63x oil objective lens, with numerical aperture of 1.4 and spatial resolution of 0.15 μm/pixel (1024 × 1024 pixels). A 488 nm wavelength laser was used to excite the SC samples. Fluorescent emissions were recorded in the range 500 to 540 nm. Perpendicular cross sectional fluorescent intensity profiles (spanning ±2.5 μm either side of the intercellular interface) across *n* = 2000 intercellular regions of neighboring corneocytes (*n* = 2 samples; *n* = 10 junctions per sample; *n* = 100 cross-sections per junction for each UV treatment condition) were extracted from the images. Individual profiles for each UV dosage condition were normalized by peak pixel intensity, then averaged. Intercellular junctions were considered to be located along the coinciding perimeters of adjacent corneocytes(42).

### 2.5 Statistical analysis

All statistical analyses were performed using R (version 3.4.2). A 1-way ANOVA was used to test for statistical significance in Figs. 1 and 2, where each UV condition (UVA, UVB, and UVC) was compared to the control. Levene’s and Shapiro-Wilk’s tests were respectively used to determine equality of variances and normality. Results in Fig. 1C (controls), Fig. 2A (UVA 10 and 200 J/cm^2^), Fig. 2B (UVB 10 J/cm^2^), Fig. 2C (UVB 800 J/cm^2^, UVC 800 J/cm^2^), and Fig. 2D (UVB 100 J/cm^2^, UVC 800 J/cm^2^) were found to exhibit non-normal distributions, but equal variances. Here a Kruskal-Wallis analysis was performed. Results in Fig. 1D (UVB), and Fig. 2B–D (UVB) were found to exhibit normal distributions, but unequal variances. Here a 1-way ANOVA with Welch correction was performed. Post-hoc analyses were performed if statistical significance levels below 5% were established. In the figures, * denotes *p* ≤ 0.05, ** denotes *p* ≤ 0.01, and *** denotes p ≤ 0.001.

**Figure 1.**
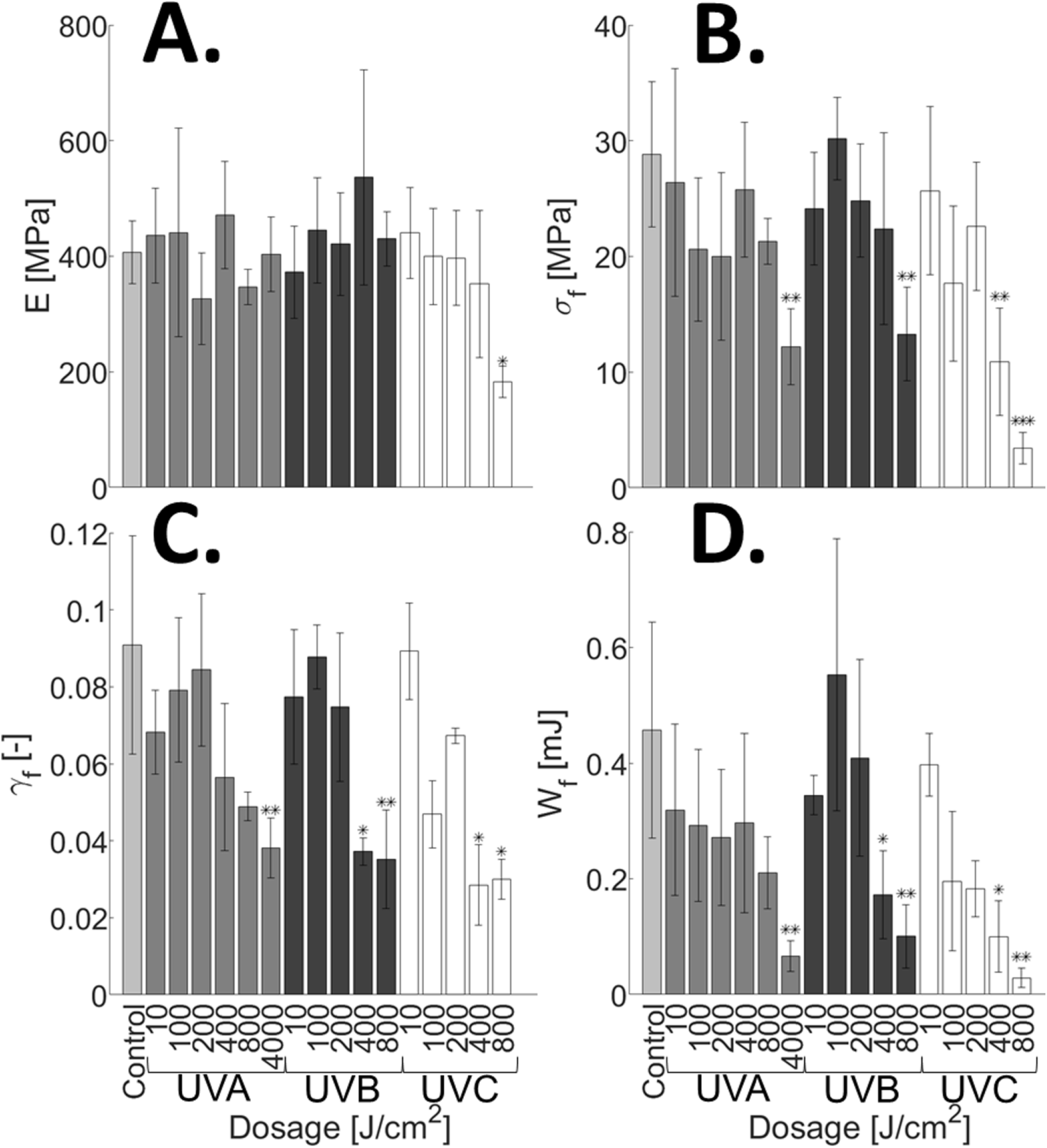
UV induces changes in the mechanical properties of SC equilibrated to 25% RH. Average (A) elastic modulus, *E*, (B) fracture stress, *σ_f_*, (C) fracture strain, *γ_f_*, and (D) work of fracture, *W_f_* for unirradiated controls (Control; light grey), UVA irradiated samples (dosage range: 10 - 4000 J/cm^2^; medium grey), UVB irradiated samples (dosage range: 10 - 800 J/cm^2^; dark grey), and UVC irradiated samples (dosage range: 10 - 800 J/cm^2^; white). Bars denote average values of 3 ≤ *n* ≤ 9 individual sample measurements for each range and dosage condition. Error bars denote standard deviations. [2 column]

**Figure 2.**
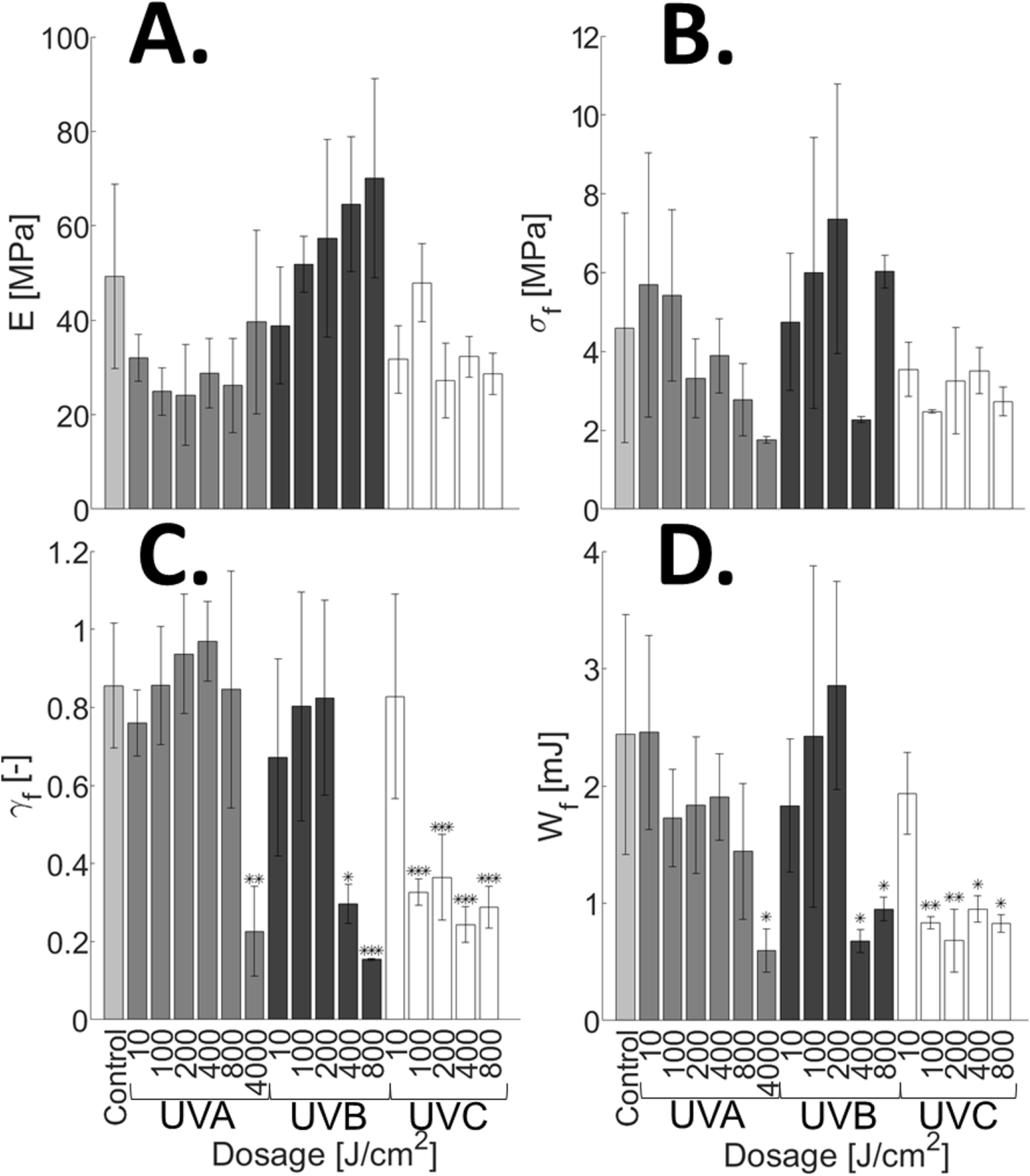
UV induces changes in the mechanical properties of SC equilibrated to 100% RH. Average (A) elastic modulus, *E*, (B) fracture stress, *σ_f_*, (C) fracture strain, *γ_f_*, and (D) work of fracture, *W_f_* for unirradiated controls (Control; light grey), UVA irradiated samples (dosage range: 10 - 4000 J/cm^2^; medium grey), UVB irradiated samples (dosage range: 10 - 800 J/cm^2^; dark grey), and UVC irradiated samples (dosage range: 10 - 800 J/cm^2^; white). Bars denote average values of 3 ≤ *n* ≤ 7 individual sample measurements for each range and dosage condition. Error bars denote standard deviations. [2 column]

## 3. RESULTS AND DISCUSSION

### 3.1 Changes in SC mechanical properties with UV irradiation

Changes in the mechanical integrity of human SC with UV light range and dosage are first assessed. Isolated human SC samples are irradiated with incident dosages of narrowband UVA (365 nm), UVB (302 nm) or UVC (265 nm) light ranging between 0 (Control) and 4000 J/cm^2^; the latter dosage being equivalent to approximately 8 continual days of full-spectrum solar UVA (or 361 continual days solely at 365 nm) (ASTM G173-03, 2012). UV therapy treatments for various skin diseases including vitiligo, pruitus, inflammatory dermatoses, and scleroderma can have cumulative dosages of up to 485 J/cm^2^ for narrowband UVB (311 nm)(44–46) and 2000 J/cm^2^ for broadband UVA (340 to 400 nm)(47–49). Figs. 1A–D respectively show the average (3 ≤ *n* ≤ 9 samples for each UV dosage) elastic modulus, *E*, fracture stress,*σ_f_*, fracture strain, *γ_f_*, and work of fracture, *W_f_*, of irradiated samples equilibrated for 24 hr. to 25% relative humidity (RH) prior to mechanical testing. Fig. 1A highlights that no significant change in elastic modulus occurs with either UVA or UVB irradiation, however a UVC dosage of 800 J/cm^2^ induces a significant decrease in tissue stiffness, relative to controls. Fig. 1B shows that samples undergo no significant reduction in fracture stress for UVA dosages less than 4000 J/cm^2^, UVB dosages less than 800 J/cm^2^, and UVC dosages less than 400 J/cm^2^. However, dosages at or above these levels cause statistically significant decreases. Similarly, Figs. 1C and D show that significant decreases in both the fracture strain and work of fracture occur with UVA dosages of 4000 J/cm^2^, and both UVB and UVC dosages equal to or greater than 400 J/cm^2^.

Figs. 2A–D show complementary mechanical results for SC samples (3 ≤ *n* ≤ 7 samples for each dosage) equilibrated for 24 hr. to 100% RH prior to mechanical testing. In contrast to the 25% RH results in Fig. 1, Figs. 2A and B highlight that no significant change in either the elastic modulus or fracture stress occurs for any UV treatment. Reductions in sample fracture strain and work of fracture however occur with UVA and UVB dosages of 4000 J/cm^2^ and 400-800 J/cm^2^ respectively, as shown in Figs. 2C and D. This trend is similar to that observed for samples equilibrated to 25% RH (Fig. 1C and D), however the magnitude by which these parameters decrease is notably larger for the hydrated SC. Moreover, UVC dosages of 100 J/cm^2^ or greater also induce statistically significant decreases. The lack of statistically significant changes in elastic modulus (Fig. 1A and 2A) with UVB irradiation, along with observed decreases in fracture strain with increasing UVB dosage (Fig. 1C and 2C) are consistent with previous studies(31).

Results in Fig. 1 and 2 appear to highlight that for equivalent incident dosages, UVC light induces the greatest loss in SC mechanical integrity, followed by UVB, then UVA. However, the incident dosage does not equate with the physical energy absorbed by the SC tissue (50, 51). UV light is absorbed, reflected, and transmitted through the SC; the degree with which each of these occurs is wavelength dependent. Prior studies have shown that the percentage of incident UVA, UVB, and UVC light absorbed by the SC is 25, 48, and 71%, respectively(50). Table S1 shows the energy physically absorbed by the SC for each UV range and incident dosage. We further scrutinize our data to establish whether a relationship exists between the energy absorbed by the SC, and the associated degradation in its mechanical integrity.

### 3.2 Scaling of absorbed UV dosage with work of fracture

Figs. 3 and 4 respectively show the work of fracture plotted against the physical energy absorbed by individual SC samples, when equilibrated to 25 and 100% RH. Both figures reveal that irrespective of the UV range used, increases in the energy absorbed by the tissue decrease the work of fracture. For each figure, a relationship between the average work of fracture, *W_f_*, and the absorbed dosage, *ϕ*, is established by fitting the data with a nonlinear least-squares regression of the form;

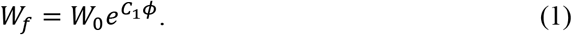

**Figure 3.**
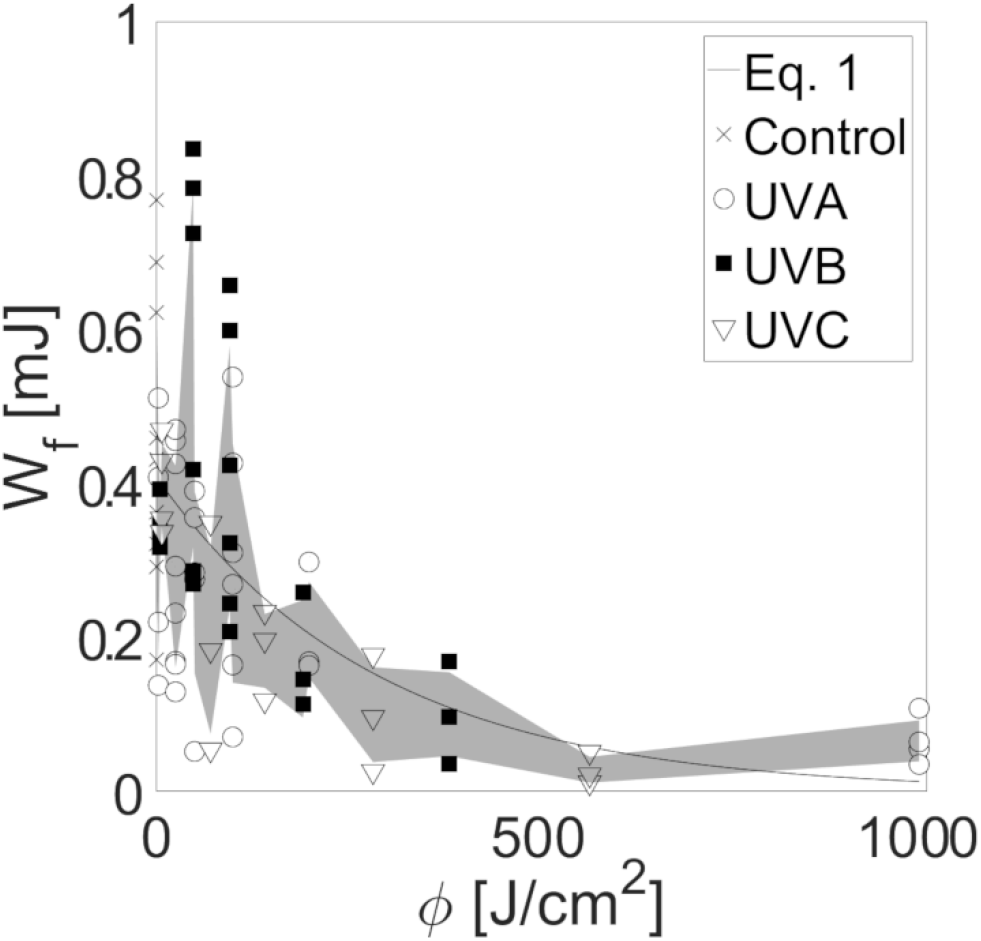
Change in the work of fracture with UV energy absorbed by SC equilibrated to 25% RH. Work of fracture, *W_f_*, of individual SC samples plotted against absorbed UV dosage, ϕ, for controls (cross), and samples irradiated with UVA (open circle), UVB (filled square), and UVC (open inverted triangle). An exponential least-squares best fit is represented by the solid black curve (Eq. 1, *W*_0_ = 0.395 mJ, *C*_1_ = −3.61 × 10^−3^). The shaded region indicates the standard deviation of the data points about the mean work of fracture at each dosage. [single column]

**Figure 4.**
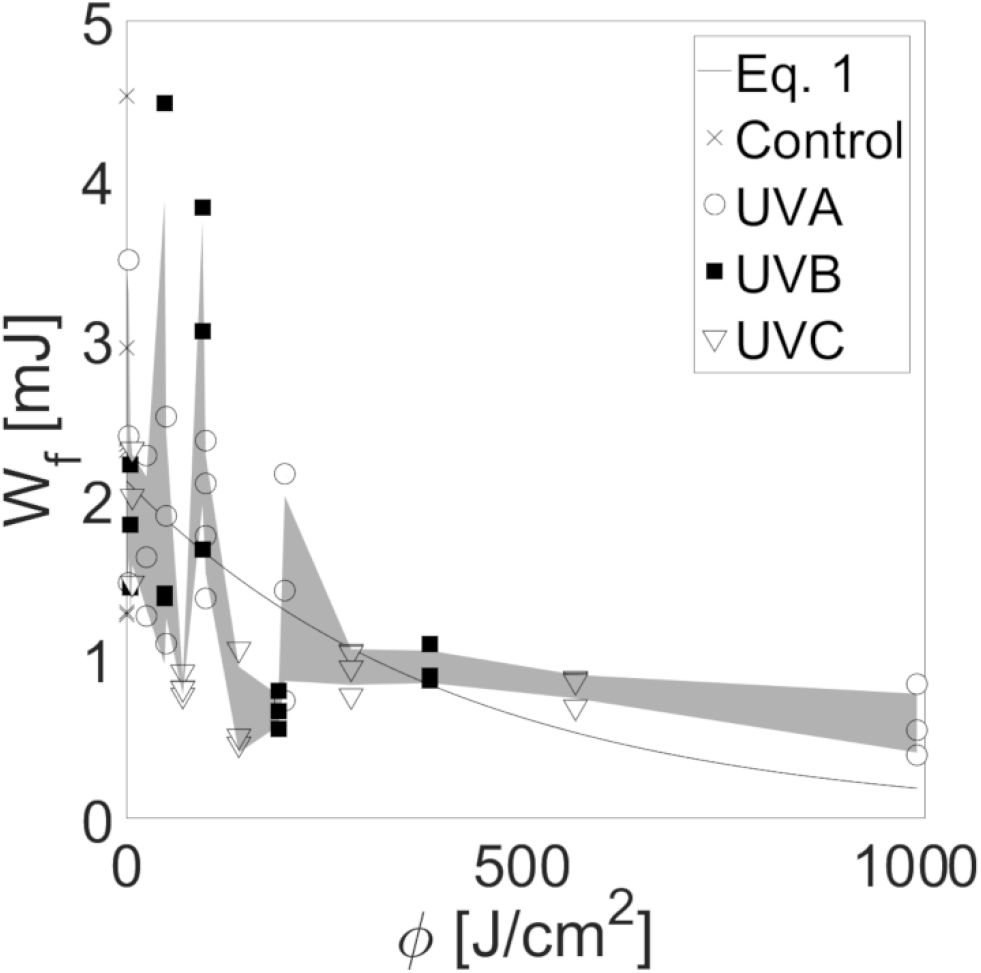
Change in the work of fracture with UV energy absorbed by SC equilibrated to 100% RH. Work of fracture, *W_f_*, of individual SC samples plotted against absorbed UV dosage, ϕ, for controls (cross), and samples irradiated with UVA (open circle), UVB (filled square), and UVC (open inverted triangle). An exponential least squares best fit is represented by the solid black curve (Eq. 1, *W*_0_ = 2.01 mJ, *C*_1_ = −1.73 × 10^−3^). The shaded region indicates the standard deviation of the data points about the mean work of fracture at each dosage. [single column]

Best fits were made with power-law, exponential, and linear models. The exponential model in each case provides the strongest correlation. The grey shaded region in each figure corresponds to the standard deviation about the average work of fracture at each dosage. For tissue equilibrated to 25% RH, *W*_0_ = 0.395 mJ and *C*_1_ = −3.61 × 10^−3^. A p-value of *p* = 5 × 10^−4^ for the goodness of fit supports the notion that absorbed dosage acts as a predictor of tissue mechanical integrity, with an R-squared value of 0.69. Likewise, for tissue equilibrated to 100% RH, *W*_0_ = 2.01 mJ and *C*_1_ = −1.73 × 10^−3^. A p-value of *p* = 6× 10^−3^ is also established for the goodness of fit, with an R-squared value of 0.48. These results indicate that irrespective of the UV range used, the dosage absorbed by the tissue appears to be primary driving force behind the degradation of mechanical integrity.

### 3.3 Dispersion of intercellular desmoglein 1 with UV irradiation

While our results clearly demonstrate a loss of tissue mechanical integrity with sufficient irradiation from all UV ranges, the underlying cause of this degradation remains unclear. Previous studies have highlighted that tissue hydration is the predominant factor in altering the mechanical properties of SC(52–56). Reductions in water content are associated with increases in tissue elastic modulus(52–56), and notable decreases in the ability of SC to plastically deform prior to rupture(53, 55). While reductions in plastic deformability of SC equilibrated to 100% RH do occur with UV irradiation, as demonstrated in Fig. 5, associated increases in elastic modulus are absent (Figs. 1A and 2A). Therefore, a structural change is more likely to be the primary cause of the observed SC mechanical degradation. Alterations to the corneodesmosome junctions is an attractive explanation due to its role as a major structural protein that provides intercellular cohesion(57, 58). The impact of UV irradiation on changes to the presence and location of desmoglein 1 (Dsg 1), a key component of corneodesmosomes, is consequently investigated using immunostaining.

**Figure 5.**
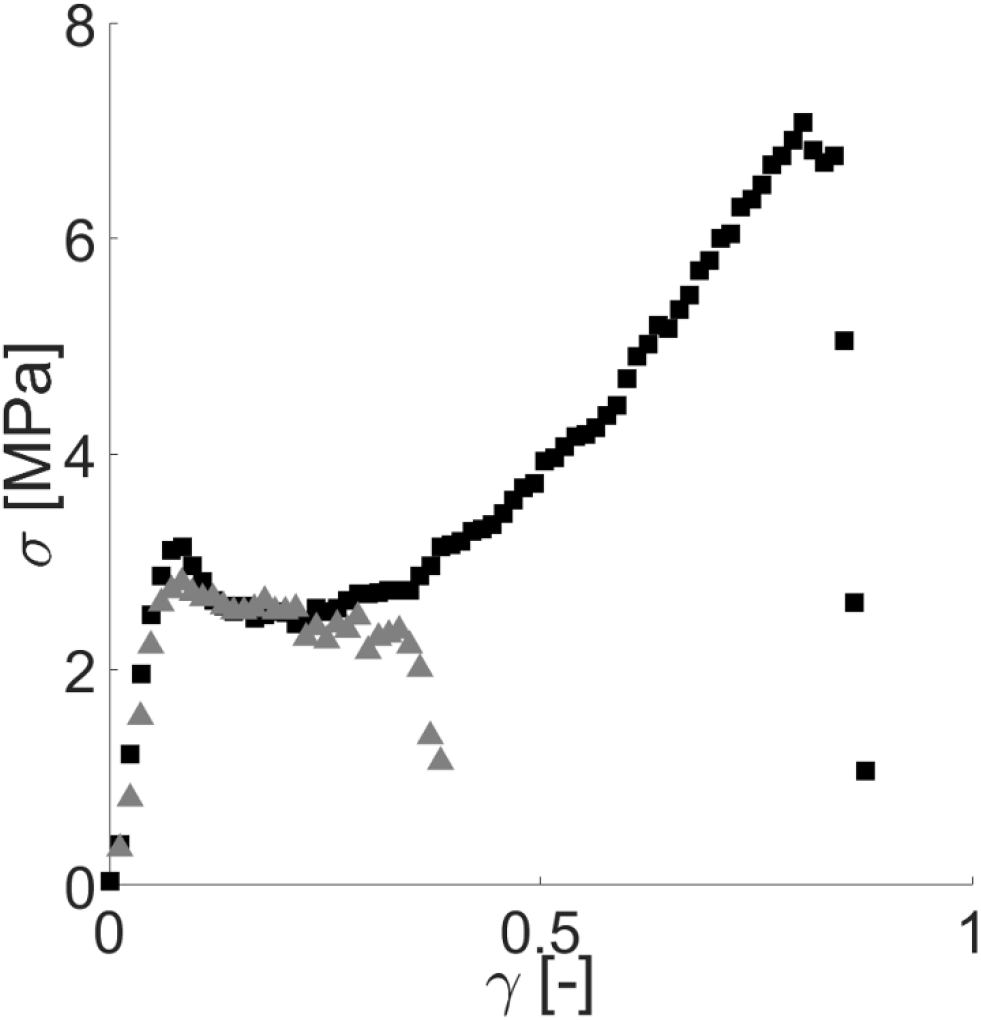
Loss of SC plastic deformability with UV irradiation. Representative stress-strain plots of an unirradiated SC sample (filled squares) and a sample irradiated with 400 J/cm^2^ of incident UVB light (filled triangle). Both samples are equilibrated to 100% RH. [single column]

Fig. 6 shows the change in distribution of fluorescently tagged Dsg 1 in SC samples irradiated with increasing UVB exposure. Control SC samples exposed only to ambient UV in Fig. 6A and B exhibit commonly observed Dsg 1 distributions, characterized by small, distinct puncta surrounding the periphery of corneocytes(40, 41, 59). Incident UVB dosages of 100 J/cm^2^ (Fig. 6C and D) and 200 J/cm^2^ (Fig. 6E and F) display similar distributions. However Fig. 6G–J show that with incident UVB dosages of 400 J/cm^2^ or greater, Dsg 1 lose their distinctive puncta morphology, and become more dispersed. Distribution changes in Dsg 1 with UVB exposure has been cited previously with organotypic cultures, but not to our knowledge with *ex vivo* human SC(59).

**Figure 6.**
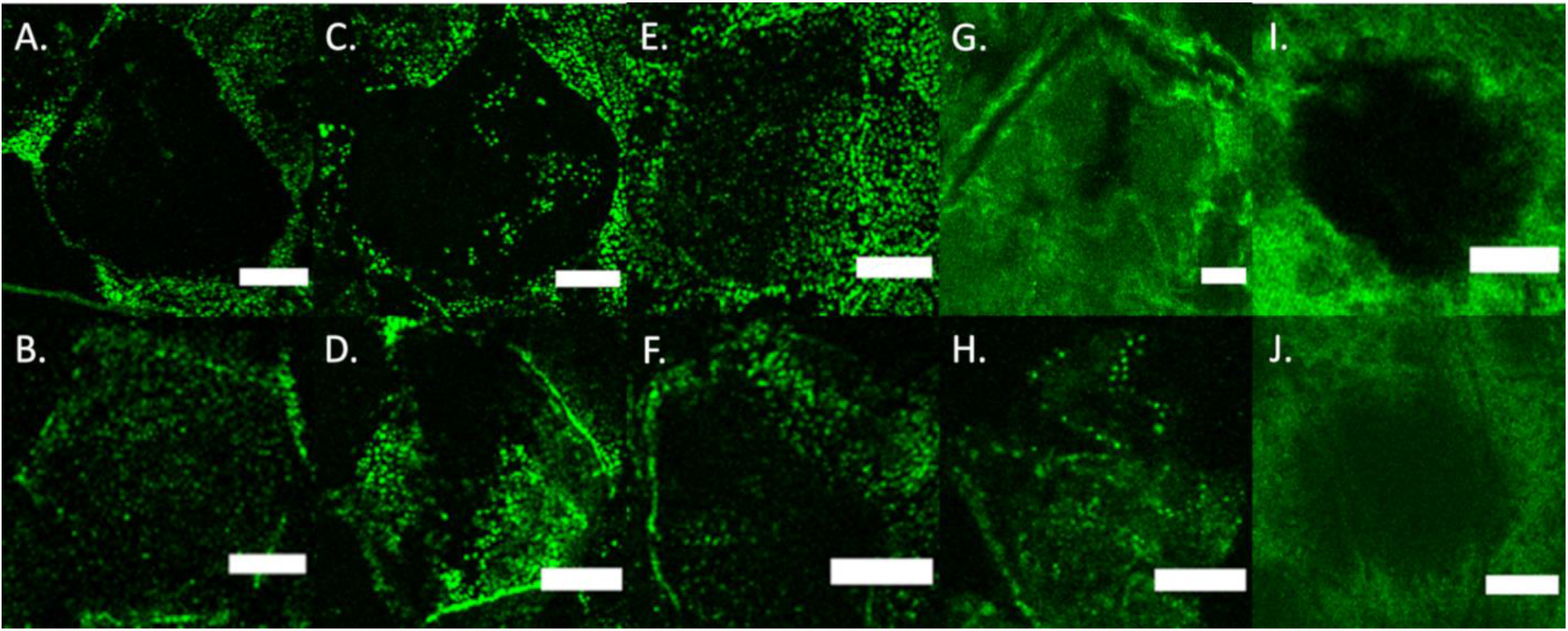
Confocal images showing changes in fluorescently tagged desmoglein (Dsg 1) distributions with increasing UVB dosage. Dsg 1 fluorescent images (from *n*=2 individual samples per condition) of control samples (A,B) and samples irradiated with an incident dosage of 100 J/cm^2^ (C,D), 200 J/cm^2^ (E,F), 400 J/cm^2^ (G,H), and 800 J/cm^2^ UVB (I,J). (*Scale bar*) 10 μm. [2 column]

The progressive dispersion of Dsg 1 is further quantified across all UV types (UVA, UVB and UVC) and dosages (0-800 J/cm^2^) by characterizing the average fluorescent normalized pixel intensity (*NPI*) profiles perpendicularly across intercellular junctions. Average *NPI* profiles for each UV range and dosage are established from *n* = 2000 intercellular junction profiles (*n*=2 samples; *n*=10 junctions per sample; *n*=100 cross-sections per junction). Profiles from UVB irradiated samples are displayed in Fig. 7A–E. UVA and UVC irradiated profiles are provided in Fig. S1A–D and E–H respectively in the Supporting Material. All cross-sectional profiles, irrespective of UV type and dosage, exhibit peak *NPI* values at or near the center of the cross-sectional profile, coincident with the cell-cell interface. However, while intensity profiles exhibit a narrow peak at the cell-cell interface for UVB dosages of 200 J/cm^2^ or less (Fig. 7A–C), consistent with dense localized puncta, greater dosages (Fig. 7D and E) exhibit notably wider peaks, indicative of dispersion. UVA and UVC irradiated samples also exhibit this trend, with profile peaks widening with UVA dosages greater than or equal to 800 J/cm^2^ (Fig. S1D) and UVC dosages greater than or equal to 200 J/cm^2^ (Fig. S1F–H). These results indicate that increasing dosages result in increased dispersion of desmoglein for all UV ranges. To better understand this dispersion, *NPI* profiles are fitted with a double gaussian function of the form,

**Figure 7.**
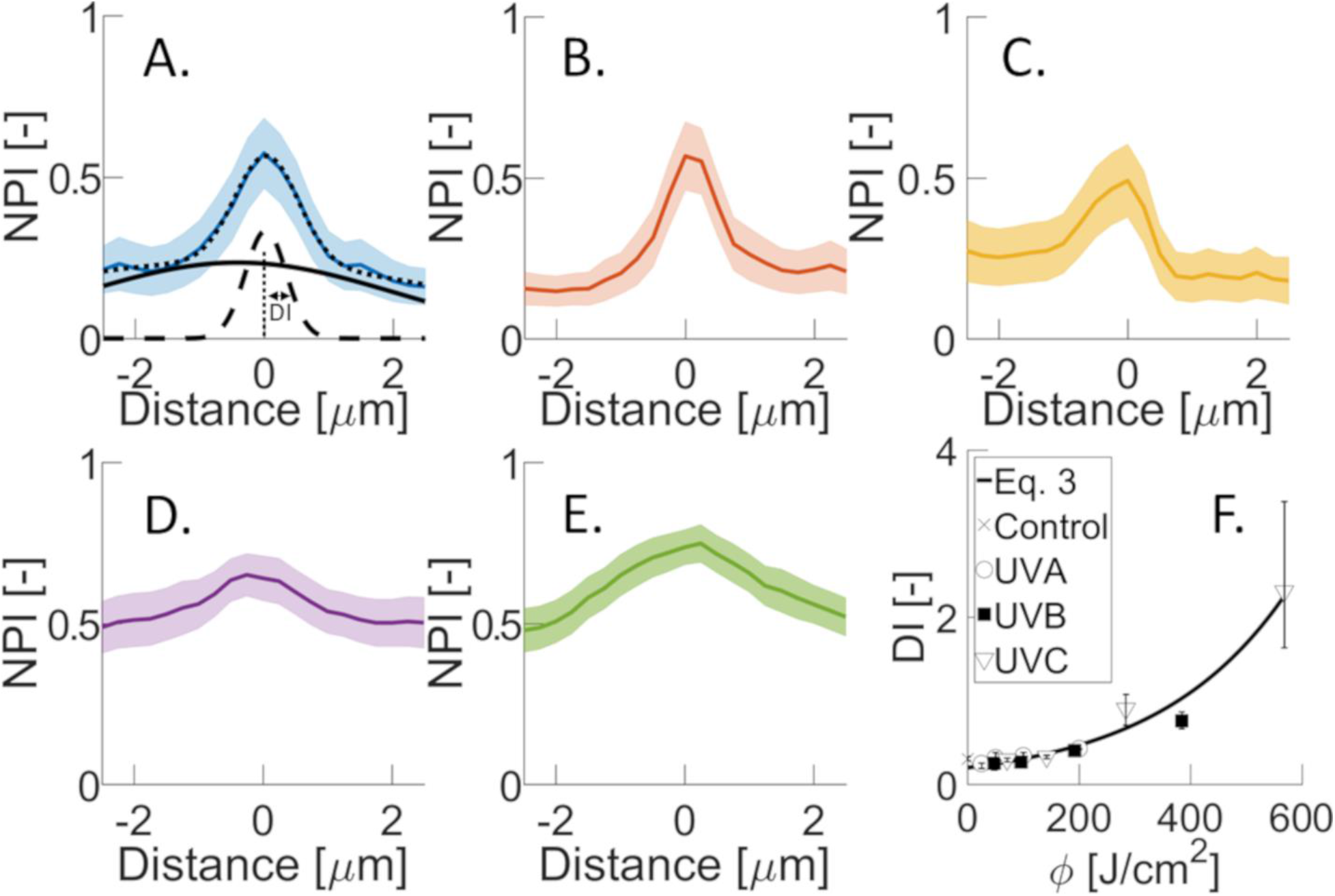
Changes in averaged normalized pixel intensity (*NPI*) profiles across intercellular corneocyte junctions with increasing UVB dosage. (A) control samples (blue line), (B) 100 J/cm^2^ incident dosage (red line), (C) 200 J/cm^2^ incident dosage (yellow line), (D) 400 J/cm^2^ incident dosage (purple line), and (E) 800 J/cm^2^ UVB incident dosage (green line). Shaded regions correspond to the standard error about the mean curve. Eq. 2 is used to fit average *NPI* profiles. The black dotted line denotes the best fit to the average control *NPI* profile in panel A. The two individual composite gaussian distributions of Eq. 2 are also represented in this panel by black solid- and dashed-lines. The inner gaussian (black dashed-line) is used to establish the dispersion index (*DI*), equivalent to the standard deviation of the distribution. (F) Change in dispersion index, *DI*, with UVA (open circle), UVB (filled square), and UVC (open inverted triangle) energy absorbed by SC. An exponential least-squares best fit to all the data points is represented by the solid black curve. Error bars denote standard deviations. [2 column]

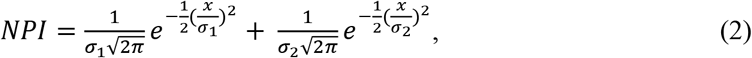

where *σ_i_* is the standard deviation and x denotes the lateral position across the averaged *NPI* profile (−2.5 ≤ *x* ≤ 2.5 μm). The best-fit model to the control treatment profile is shown as the dotted-line in Fig. 7A. The inner and outer gaussian components of Eq. 2 are also plotted in Fig. 7A as dashed and solid curves, respectively. We employ the standard deviation of the inner gaussian function to characterize the dispersion of desmoglein, which we denote the dispersion index, *DI*. Fig. 7F plots *DI* against the absorbed UV dosage, *ϕ*, irrespective of UV type. Similar to Fig. 3 and 4, an exponential function of the form,

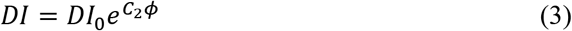

is found to provide the best fit to the data. For the regression, *DI*_0_ = 0.199 and *C*_2_ = 4.27 × 10^−3^. A p-value of *p* = 7 × 10^−4^ for the goodness of fit supports the absorbed dosage acting as a predictor of Dsg 1 distribution, with an R-squared value of 0.96.

Changes in both the average work of fracture (Fig. 3 and 4), *W_f_*, and structural dispersion index (Fig. 7F), *DI*, with absorbed UV dosage, *φ*, exhibit exponential relationships. Combining Eq. 1 and 3, we establish a power law expression relating structural and mechanical parameters of the form,

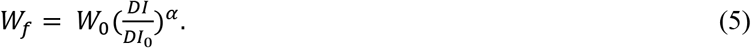

where *α* = *C*_1_/*C*_2_. For SC samples equilibrated to 25% RH, *α* = −0.845. This structure-mechanical relationship highlights that the mechanical integrity of the SC tissue scales with near inverse proportionality to the dispersion of intercellular junction proteins away from intercellular interfaces. We restrict this model to tissue equilibrated to dry (25% RH conditions) conditions because all UV irradiation protocols were performed on dry tissue samples to minimize surface, unbound, and bound water(60, 61) altering the absorption, reflection, and transmission of UV(51, 62–64). Moreover, SC equilibrated to a lower relative humidity is more physiologically consistent in-vivo SC tissue (20-40% RH indoors(65); 30-80% RH outdoors(66)).

## 4. Conclusion

In this article, we quantify the impact of UV range and dosage on the mechanical and structural degradation of isolated human stratum corneum (SC). For equivalent incident dosages, UVC light causes the greatest loss in mechanical integrity of the tissue, followed by UVB and UVA. This is due to the greater radiation energy UVC. However, when discounting reflected and transmitted components of the incident light, a scaling law relationship between the energy absorbed and the work of SC fracture emerges. This relationship highlights that no one UV range is more damaging than another. Rather, the amount of energy absorbed governs the loss of tissue mechanical integrity. Further structural studies reveal that desmoglein 1 (Dsg 1), a major component of corneodesmosomes, becomes dispersed with UV exposure away from intercellular sites. Upon relating the mechanical and structural models, a near inverse scaling law is revealed. This suggests that a simple immunostaining assay could therefore be employed to quantify UV induced tissue damage.

We anticipate that UV light induced increases in protease activity could contribute to the observed structural changes in Dsg 1 within corneodesmosomes, resulting in mechanical degradation. Stratum corneum serine proteases that hydrolyze corneodesmosome components such as Dsg 1 exhibit increased activity with UVB exposure in cultured epidermal keratinocytes(67). While reports have shown that the enzyme activity of these proteases can be modulated *ex vivo* with factors such as hydration(68) and inhibitors(69, 70), future studies should determine if similar increases in enzyme activity with UV irradiation are seen in *ex vivo* SC. In addition, the scaling relationships established in this study should be further tested. The validity of using absorbed energy and structural protein dispersion as a predictor of biomechanical photoaging should be assessed over a wider range of the electromagnetic spectrum and in deeper skin layers, perhaps through biomechanical testing and immunostaining of progressively thicker skin specimens that retain epidermal, then dermal tissue.

## AUTHOR CONTRIBUTION

G.K.G. designed the research. Z.W.L. performed the experiments and analyzed the data. Both authors wrote the article.

## ACKNOWLEDGEMENTS

This material is based upon work supported by the National Science Foundation under Grant No. 1653071. The authors state no conflict of interest.

## Abbreviations

(UV): Ultraviolet
(SC): Stratum Corneum
(RH): Relative Humidity
(Dsg 1): Desmoglein 1

